# Divergent selection causes whole genome differentiation without physical linkage among the targets in *Spodoptera frugiperda* (Noctuidae)

**DOI:** 10.1101/452870

**Authors:** Kiwoong Nam, Sandra Nhim, Stéphanie Robin, Anthony Bretaudeau, Nicolas Nègre, Emmanuelle d’Alençon

## Abstract

The process of speciation involves whole genome differentiation by overcoming gene flow between diverging populations. We have ample knowledge which evolutionary forces may cause genomic differentiation, and several speciation models have been proposed to explain the transition from genetic to genomic differentiation. However, it is still unclear what are critical conditions enabling genomic differentiation in nature. The Fall armyworm, *Spodoptera frugiperda*, is observed as two sympatric strains that have different host-plant ranges, suggesting the possibility of ecological divergent selection. In our previous study, we observed that these two strains show genetic differentiation across the whole genome with an unprecedentedly low extent, suggesting the possibility that whole genome sequences started to be differentiated between the strains. In this study, we analyzed whole genome sequences from these two strains from Mississippi to identify critical evolutionary factors for genomic differentiation. The genomic Fst is low (0.017) while 91.3% of 10kb windows have Fst greater than 0, suggesting genome-wide differentiation with a low extent. We identified nearly 400 outliers of genetic differentiation between strains, and found that physical linkage among these outliers is not a primary cause of genomic differentiation. Fst is not significantly correlated with gene density, a proxy for the strength of selection, suggesting that a genomic reduction in migration rate dominates the extent of local genetic differentiation. Our analyses reveal that divergent selection alone is sufficient to generate genomic differentiation, and any following diversifying factors may increase the level of genetic differentiation between diverging strains in the process of speciation.

## INTRODUCTION

Speciation processes inherently involve genomic differentiation by reproductive barriers, generated through collective or sequential actions of evolutionary forces(Wu 2001). However, gene flow impedes the process of speciation because recombination in hybrids homogenizes sequences between populations(Felsenstein 1981). An exceptional condition is, therefore, necessary to overcome the homogenizing effect of gene flow (reviewed in (Bolnick and Fitzpatrick 2007)). Accumulating empirical reports show that speciation indeed occurs in the presence of gene flow(Nosil 2008), implying that the homogenizing effect of recombination can be effectively overcome. One of the key issues to understand the speciation process is how the homogenizing effect of recombination is overcome throughout whole genomes(Feder, Egan, et al. 2012).

Divergent selection is one of the main players during the process of speciation. If selection is sufficiently strong (*i.e*, *s* > *m*(Flaxman et al. 2014) or *s* > *r*(Barton 1979), where *s*, *m*, and *r* are selection coefficient, migration rate, and recombination rate, respectively), the effect of selection dominates that of gene flow and recombination, thus genomic differentiation may not be hampered by gene flow. If selection is weak (*s* < *m* and *s* < *r*), other conditions are necessary for genomic differentiation. Physical linkage among the targets might be responsible for genomic differentiation, as selective sweeps(Smith and Haigh 1974) increase in the level of genetic differentiation at sites physically linked to the targets of divergent selection. For example, if divergent selection targets a large number of loci, then the average physical distance from a neutral locus to the targets decreases, thus whole genome sequences can be differentiated by the concerted actions of divergent selection(Barton and Bengtsson 1986). In another speciation model, termed divergence hitchhiking, if a locus is targeted by strong divergent selection, then the effective rate of migration is reduced in this region, and following events of divergent selection targeting sequences within this region may generate a long stretch of differentiated DNA (up to several Mb)(Via and West 2008; Via 2012). Population-specific chromosomal rearrangements can also contribute to the process of speciation because recombination is inhibited in hybrids(Noor et al. 2001; Rieseberg 2001; Butlin 2005; Kirkpatrick and Barton 2006), and physical linkage between targets of divergent selection and the loci with a chromosomal rearrangement may create long genomic regions with differentiation(Feder, Nosil, and Flaxman 2014). Whole genome sequences may be differentiated without physical linkage among targets of selection as well. According to the genome hitchhiking model, if divergent selection targets many loci, then genome-wide migration rate is effectively reduced, and whole genome sequences can be differentiated(Feder and Nosil 2010; Feder, Gejji, et al. 2012).

If the number of targeted loci is sufficiently high, genomic differentiation may occur rapidly. The loci targeted by population-specific divergent selection may have correlated allele frequencies, and corresponding linkage disequilibrium among targets will be then generated(Barton 2010; Flaxman et al. 2014; Schilling et al. 2018). Theoretical studies(Barton 2010; Flaxman et al. 2014) show that if the number of targets is higher than a certain threshold, targeted loci have a synergistic effect in increasing linkage disequilibrium among targets, thus genomic differentiation is consequently accelerated. This non-linear dynamics of genomic differentiation according to the number of occurred selection events were termed genome-wide congealing(Feder, Nosil, Wacholder, et al. 2014). It should be noted that any diversifying factors, including divergent selection, background selection, and assortative mating(Kopp et al. 2017), may contribute to genome-wide congealing. Thus, the critical question of how genomic differentiation occurs in the presence of gene flow is the condition for the transition to the phase of genome-wide congealing. For example, divergence hitchhiking may provide a condition for genome-wide congealing(Feder, Egan, et al. 2012). Alternatively, genome-wide reduction in migration rate (genome hitchhiking) or chromosomal rearrangement may contribute to this condition as well.

Divergence hitchhiking model has been supported by pea aphids(Via 2012), stickleback(Marques et al. 2016), and poplar(Ma et al. 2018). However, as Feder and Nosil demonstrated(Feder and Nosil 2010), long differentiated sequences can be observed only from a specific condition, when effective population size (*Ne*) and migration rate are low (*Ne* = 1,000, *m* = 0.001), and selection is very strong (*s* = 0.5). Isolation by adaptation is a positive correlation between a genetic difference and adaptive divergence(Nosil et al. 2008; Nosil et al. 2009), and this observation has been presented as a support for genome hitchhiking, which indeed causes isolation by adaptation(Feder, Egan, et al. 2012). However, it is still unclear whether genome hitchhiking initiates or reinforces genetic differentiation in cases of isolation by adaptation.

The Fall armyworm, *Spodoptera frugiperda*, (Lepidoptera, Noctuidae) is a pest species observed as two sympatric strains, corn strain (sfC hereafter) and rice strain (sfR) named from their preferred host-plants, throughout North and South American continents(Pashley 1986). Based on maker-genotyping, these two strains appear to have different DNA sequences(Pashley 1986; Kergoat et al. 2012). In a wide geographical range in North America, 16% of individuals were reported to be hybrids between strains(Prowell et al. 2004), suggesting frequent gene flow. In our previous study, we observed that these two strains have a weak but significant genomic differentiation (Fst = 0.019, p < 0.005), and that the differentiated loci were distributed across the whole genome(Gouin et al. 2017). As this level of genomic differentiation is one of the lowest among reported cases, and hybrids are frequently generated(Prowell et al. 2004), these two strains an ideal system to explore critical evolutionary forces for genomic differentiation in the presence of gene flow. Whole genome differentiation between sfC and sfR might involve both premating reproductive isolation through assortative mating(Schöfl et al. 2009; Unbehend et al. 2013; Hänniger et al. 2017), or postmating reproductive isolation by ecological divergent selection, or by reduced hybrid fertility(Dumas, Legeai, et al. 2015).

In this study, we aim at identifying evolutionary forces that are responsible for genomic differentiation between sfC and sfR at the very initial stage of the speciation process. Using resequencing data generated in our previous study(Gouin et al. 2017), we test the role of several evolutionary forces in genomic differentiation, including chromosomal rearrangements, physical linkages among targeted loci, and genomic reduction in migration rate. The results presented here allow us to identify critical evolutionary factors that enable the genomic differentiation between strains in *S. frugiperda*.

## RESULTS

It is important to have a contiguous reference genome assembly to accurately detect signatures of genome divergence. The reference genome assemblies for sfC and sfR generated from our previous study contain 41,577 and 29,127 scaffolds, respectively(Gouin et al. 2017) (Table 1). We performed *de novo* genome assembly from Pac-bio reads (27.5X and 33.1X coverages for sfC and sfR, respectively) to improve the reference genome sequences. Errors in these reads were corrected by Illumina assemblies, which were generated from the reads used in our previous study(Gouin et al. 2017). The Pac-bio reads were assembled using SmartDenovo(Ruan 2017), and scaffolding was performed using Illumina paired-ends and mate-pairs used in our previous study. The resulting assemblies are now closer to the expected genome sizes, 396±3Mb, estimated by flow cytometry(Gouin et al. 2017) (Table 1). Moreover, the contiguity is also significantly improved, as N50 is 900kb and 1,129kb for corn and rice reference genome sequences, respectively. The numbers of sequences are 1,000 and 1,054 for sfC and sfR, respectively. BUSCO analysis(Simão et al. 2015) shows that the correctness is also increased, especially for the sfC (Supplementary Table 1). The numbers of identified protein-coding genes are 21,839 and 22,026 for sfC and sfR, respectively. BUSCO analysis shows that gene annotation is also improved, especially for sfC (Supplementary Table 2).

**Table 1.** Summary statistics of genome assemblies produced in this study (New assembly) and published assembly(Gouin et al. 2017) from corn and rice strains.

|  | Corn strain |  | Rice strain |  |
| --- | --- | --- | --- | --- |
| statistics | New assembly | Gouin et al | New assembly | Gouin et al |
| Assembly size | 384,358,373 | 437,873,304 | 379,902,278 | 371,020,040 |
| number of sequences | 1,000 | 41,577 | 1,054 | 29,127 |
| Longest sequence (bp) | 5,279,935 | 943,242 | 7,849,854 | 314,108 |
| Shortest sequence (bp) | 8,866 | 888 | 10,636 | 500 |
| N50 | 900,335 | 52,781 | 1,129,192 | 28,526 |
| L50 | 124 | 1,616 | 91 | 3,761 |
| N90 | 196,225 | 3,545 | 165,330 | 6,422 |
| L90 | 450 | 18,789 | 421 | 13,881 |
| %GC | 36.3432 | 35.0770 | 36.3724 | 36.0741 |
| %N | 0.0689 | 2.5989 | 0.0006 | 0.0352 |

Resequencing data from nine female individuals from each of corn and rice strains collected in the wild(Gouin et al. 2017) were mapped against these two nuclear reference genome assemblies using bowtie2(Langmead and Salzberg 2012: 2) with very exhaustive search parameters (see methods). As one individual from rice strain has a particularly low mapping rate and an average read depth (denoted as R1,Gouin et al.(Gouin et al. 2017)) (Supplementary Figure 1), we excluded this individual from the following analysis. Variants were called using samtools mpileup(Li et al. 2009), and we performed stringent filtering by discarding all sites unless Phred variant calling score is higher than 40, *and* genotypes are determined from every single individual. The numbers of variants are 48,981,416 from 207,415,852 bp and 49,832,320 from 205,381,292 bp from the mapping against sfC and sfR reference genomes, respectively. As analyses from the resequencing data might be affected by ascertainment bias, we performed all analyses based on these two reference genomes. We present the results only from the sfC reference genomes in the main text unless mentioned specifically. The results from the sfR reference genome are shown in the supplementary information (Supplementary Figure 14-21).

The genome-wide Fst calculated between sfC and sfR is 0.017, which is comparable to our previous study (0.019)(Gouin et al. 2017). As this low level of differentiation could be caused by chance, we calculated Fst from randomized groupings with 500 replications. We observed that no randomized grouping has higher Fst than the grouping according to strains (equivalent to p < 0.002). Thus, we concluded that the genomic sequences are significantly differentiated between strains, as we did in our previous study(Gouin et al. 2017). This genomic differentiation can be either caused by a few loci with very high levels of differentiation or by many loci with low levels of differentiation. To test these two possibilities, we calculated Fst in 10 kb window. Among total windows, 91.3% of these windows have Fst greater than 0 (Figure 1), supporting the latter explanation. The low level of genetic differentiation implies that these two strains do not experience genome-wide congealing yet.

**Figure 1.**
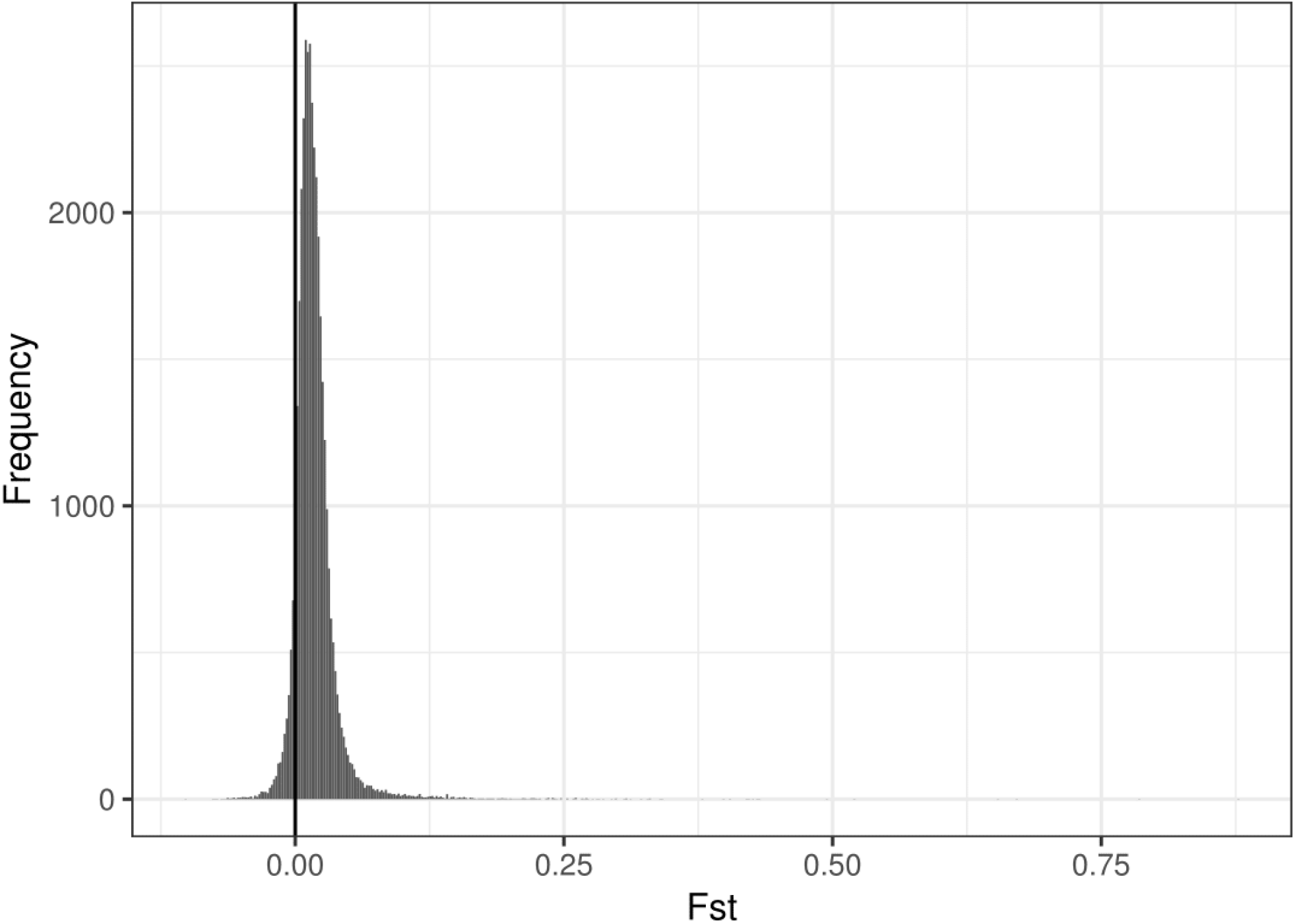
The distribution of Fst calculated in 10 kb window. The vertical line indicates Fst = 0, which means no genetic differentiation between corn and rice strains.

Genetic relationships among individuals were inferred using principal component analysis (PCA). The result shows that sfR has a higher genetic variability among individuals than sfC, and we hypothesized that sfC was derived from ancestral sfR (Figure 2a). To test this hypothesis, we reconstructed a phylogenetic tree using assembly-free approach(Fan et al. 2015) with *S. litura(Cheng et al. 2017)* as an outgroup. The resulting tree shows that sfC individuals constitute a monophyletic group, implying that the sfC was indeed derived from ancestral sfR (Figure 2b). The pattern of the phylogenetic tree is subtly different from that of PCA. The phylogenetic tree shows that sfC has monophyly, implying that the sfC individuals were derived from a single individual. However, the result from PCA does not support the single origin of sfC. This discrepancy is perhaps caused by an incomplete lineage sorting in the ancestry of sfC or by frequent gene flow between sfC and sfR. However, we cannot exclude the possibility of statistical artifacts, such as long-branch attractions(Huelsenbeck and Hillis 1993). The genetic relationship among individuals was also analyzed from ancestry coefficient(Frichot et al. 2014), and we observed that distinct origins of sfC and sfR are not supported (Supplementary Figure 2).

**Figure 2.**
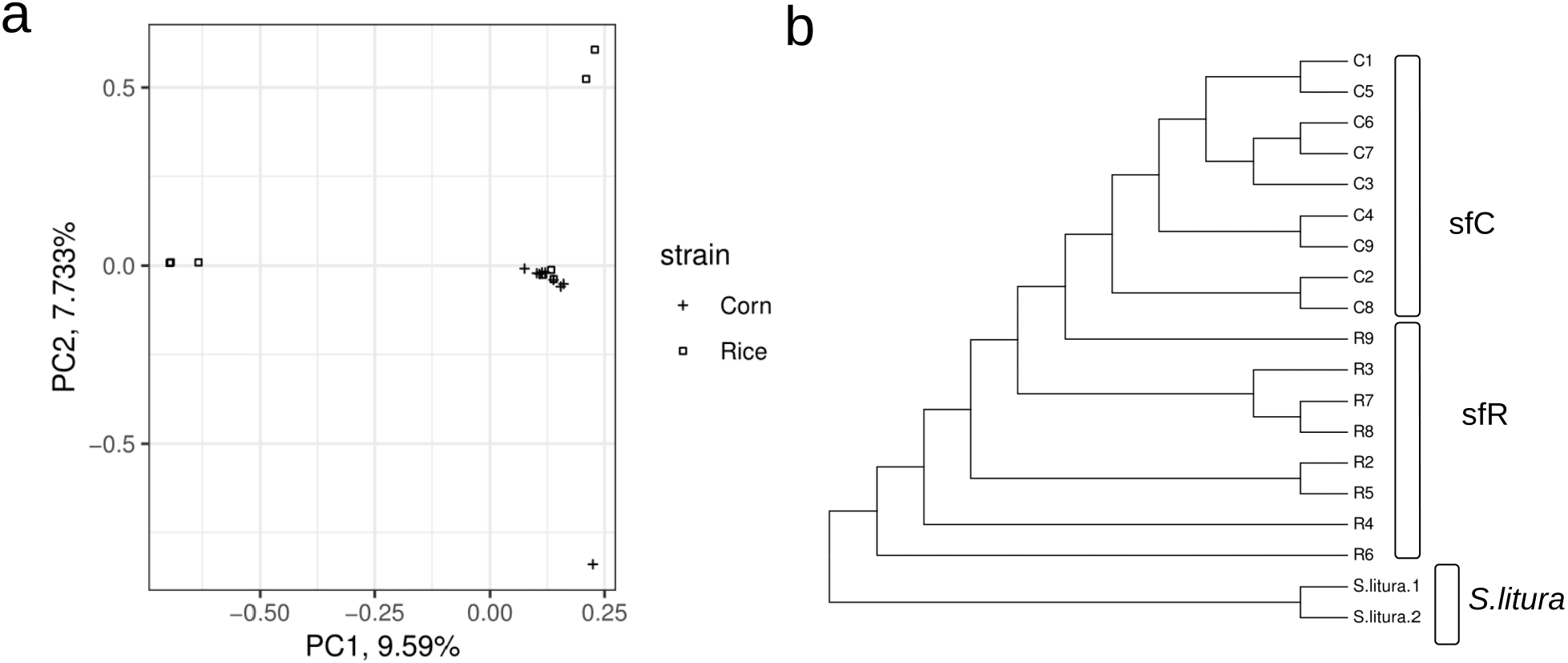
Genetic relationship between corn and rice strains. a) The result from principal component analysis. The red and blue circles represent individuals from corn and rice strains, respectively. b) Phylogenetic tree reconstructed using AAF approach.

We tested the possibility of an extreme case where both sfC and sfR have monophyly, but all identified sfR individuals except R6 on Figure 2b are F1 hybrids between sfR females and sfC males. In this case, maternally-derived mitochondrial CO1 genes used to identify strains in this study(Gouin et al. 2017) will have distinctly different sequences between R2-R9 and C1-C9, while paternally derived sequences will not show such a pattern. As all individuals analyzed in this study are females, the Z chromosomes were derived from males in the very previous generation. Thus, we tested significant genetic differentiation of Z chromosomes between sfC and sfR without R6. TPI gene is known to be linked to Z chromosomes in *S. frugiperda*(Nagoshi 2010), and we observed this gene from Contig269 by blasting. This contig is 3,688,019bp in length, and the number of variants is 201,075. The Fst calculated between sfC and sfR without R6 is 0.061, which is higher than the genomic average (0.017). We calculated Fst from randomized groupings with 500 replicates, and only four replicates have Fst higher than 0.061, corresponding p-value equal to 0.008. This result demonstrates a significant genetic differentiation of paternally derived Z chromosomes between strains that were identified by mitochondrial sequence, and we exclude the possibility of the extreme case with F1 hybrids.

We inferred changes in *Ne* from two statistics, π and Watterson’s θ. Watterson’s θ represents more recent levels of genetic diversity than π. The calculated π is 0.043 and 0.044 for sfC and sfR, respectively. The π is not significantly different between sfC and sfR (p = 0.27, permutation test with 100 randomizations). The calculated Watterson’s θ is 0.064 and 0.061 for sfC and sfR, respectively, and sfC has higher Watterson’s θ than sfR (p < 0.01). This result indicates that both sfC and sfR experienced population expansion with a greater extent in sfC, possibly due to higher fitness in sfC.

Chromosomal rearrangements specific to a single population can cause a genetic differentiation because recombination is inhibited in hybrids(Rieseberg 2001; Butlin 2005; Kirkpatrick and Barton 2006). Thus, we estimated the role of chromosomal rearrangements in genomic differentiation by identifying strain-specific chromosomal rearrangements. We identified 1,254 loci with chromosomal inversions with 1,060bp in median sequence length using BreakDancer(Chen et al. 2009). We considered that a chromosomal rearrangement is strain-specific if the difference in allele frequency is higher than an arbitrarily chosen criterion, 0.75. Fst calculated from these inversions are lower than zero (−0.063 and −0.064), meaning that the contribution of chromosomal inversion to genetic differentiation is not supported. The number of inter-scaffold rearrangement is 1,724, and only one of them has a difference in allele frequency higher than 0.75. Fst calculated from 10kb flanking sequences of the breaking points is lower than zero (−0.115 and −0.0783 at each side). Thus, we excluded the possibility that chromosomal rearrangement is a principal cause of genomic differentiation.

Then, we test the possibility that selection is responsible for genomic differentiation from outliers of genetic differentiation. We used correlated haplotype score(Fariello et al. 2013) to estimate the level of genetic differentiation between strains. If each of minimum 100 consecutive SNPs in minimum 1kb has a significantly greater haplotype score than the rest of the genome (p < 0.001), we defined this locus as an outlier. As the mapping rate of reads against highly differentiated sequences is necessarily low, the identification of outliers can be severely affected by the usage of reference genome sequences. Therefore, here we present the results from both corn and rice reference genome sequences (refC and refR, respectively). In total, 433 outliers at 170 scaffolds and 423 outliers at 148 scaffolds were identified from the mappings against refC and refR, respectively. The average length of these outliers is 4,023bp and 4,095bp for refC and refR, respectively. The longest outlier is 27,365bp and 33,110bp in length for refC and refR, respectively. These outliers occupy only small fractions of the scaffolds (1.56% and 1.82% for refC and refR, respectively). Therefore, extremely strong selective sweeps are not supported. Thus, it is unlikely that very strong selection targeting these regions causes whole genome differentiation.

We test the possibility of the divergence hitchhiking(Via 2012), a hypothesis that a strong selection creates DNA sequences with reduced local migration rate, and following selection events within this sequence generates a long stretch of DNA sequence with an elevated level of genetic differentiation. According to this speciation model, lowly differentiated sequences between highly differentiated sequences are generated by ancestral polymorphisms, rather than gene flow(Via 2012). Thus, these lowly differentiated sequences between highly differentiated sequences will show clustered ancestry maps according to the extant strains, whereas the rest of lowly differentiated sequences in the genome will not show such a clustering. From the scaffolds with the outliers, we identified lowly differentiated sequences (hapflk score < 1, Supplementary Figure 3 to see the histogram of all positions at these scaffolds), 154,163bp and 273,797bp in total size from refC and refR, respectively. Then, sNMF software was used to infer ancestry coefficients(Frichot et al. 2014). Figure 3 shows that sfC and sfR have different ancestry at outliers, while the lowly differentiated sequences within the scaffolds with outliers do not show any apparent clustering according to extant strains. Thus, divergence hitchhiking is not supported by our data.

**Figure 3.**
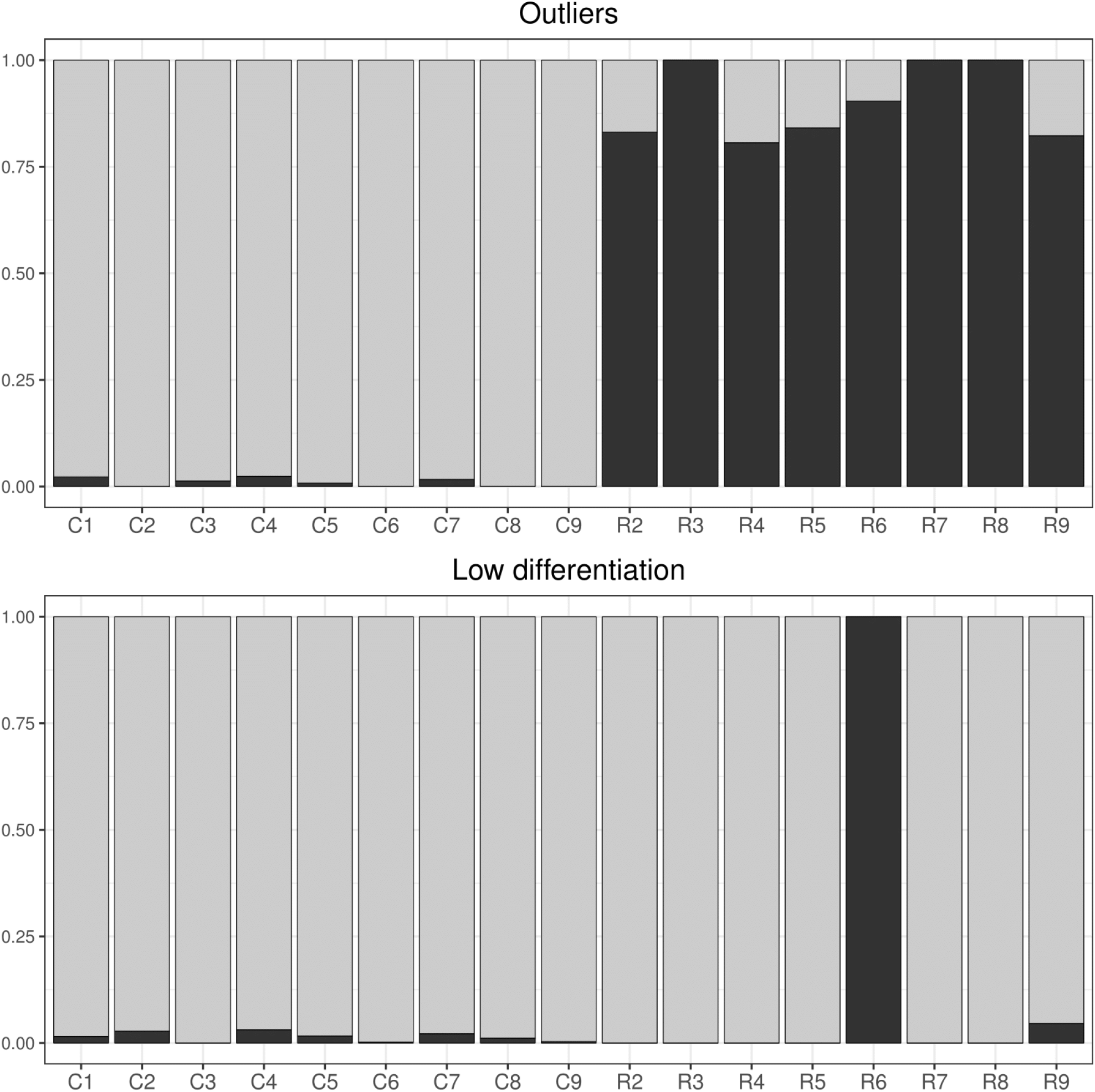
Testing the divergence hitchhiking model. Ancestry coefficient calculated from the outliers of genetic differentiation (upper) and lowly differentiated sequences (hapflk score < 1, 154,163bp in size) (bottom).

If a genetic locus is resistant against gene flow from the beginning of genetic differentiation, this sequences is expected to show a higher level of absolute genetic divergence, which can be estimated from dXY statistics(Cruickshank and Hahn 2014). We observed that four out of the 433 outliers from refC and nine out of the 423 outliers from refR have higher dXY than genomic average (FDR corrected p < 0.05) (Supplementary Figure 4, 5). We denote these outliers as genomic islands of divergence in this paper. These genomic islands of divergence contain three and four protein-coding genes from refC and refR, respectively. These genes include NPRL2 and Glutamine synthetase 2. NPRL2 is a down-regulator of TORC1 activity, and this down-regulation is essential in maintaining female fecundity during oogenesis in response to amino-acid starvation in Drosophila(Wei and Lilly 2014). Glutamine synthetase 2 is important in activating TOR pathway, which is the main regulator of cell growth in response to environmental changes to maintain fecundity in planthoppers(Jacinto and Hall 2003). This result raises the possibility that disruptive selection on female fecundity is responsible for initiating genetic differentiation between strains. The function of the other five genes is unclear. Thus, other traits might be important in initiating genomic differentiation as well.

If genetic differentiation is initiated by selection on female fecundity, mitochondrial genomes will show a higher level of absolute level of sequence divergence than nuclear genome because mitochondrial genomes are transmitted only through the maternal lineage. We performed mapping all reads against mitochondrial genomes (KM362176) and identified 371 variants from 15,230bp. The result from PCA shows that, contrary to the nuclear pattern, sfC and sfR individuals fall into two distinct groups (Figure 4a). Ancestry coefficient analysis shows that each of two strains has a distinct ancestry (Figure 4b) (see Supplementary Figure 6 to find a correlation between K and cross entropy). To generate a mitochondrial phylogenetic tree, we extracted sequences of *S. frugiperda* from mitochondrial Variant Call Format file, and we created a multiple sequence alignment together with the mitochondrial genome sequence of *S. litura* (KF701043). Then, a phylogenetic tree was reconstructed using the minimum evolution approach(Lefort et al. 2015). The tree shows that sfC and sfR are a sister group of each other(Figure 4c). This mitochondrial pattern is also observed from other studies in *S. frugiperda*(Kergoat et al. 2012; Dumas, Barbut, et al. 2015; Gouin et al. 2017). We excluded a possibility that strong linked selection on mitochondrial genomes alone causes the different phylogenetic pattern between nuclear and mitochondrial genomes because in this case the topology is expected to be unchanged while only relative lengths of ancestral branches to tips are different between nuclear and mitochondrial trees (Supplementary Figure 7). Instead, this pattern can be explained by an ancient divergence of mitochondrial genomes, which is followed by a gradual genetic differentiation of nuclear genomes.

**Figure 4.**
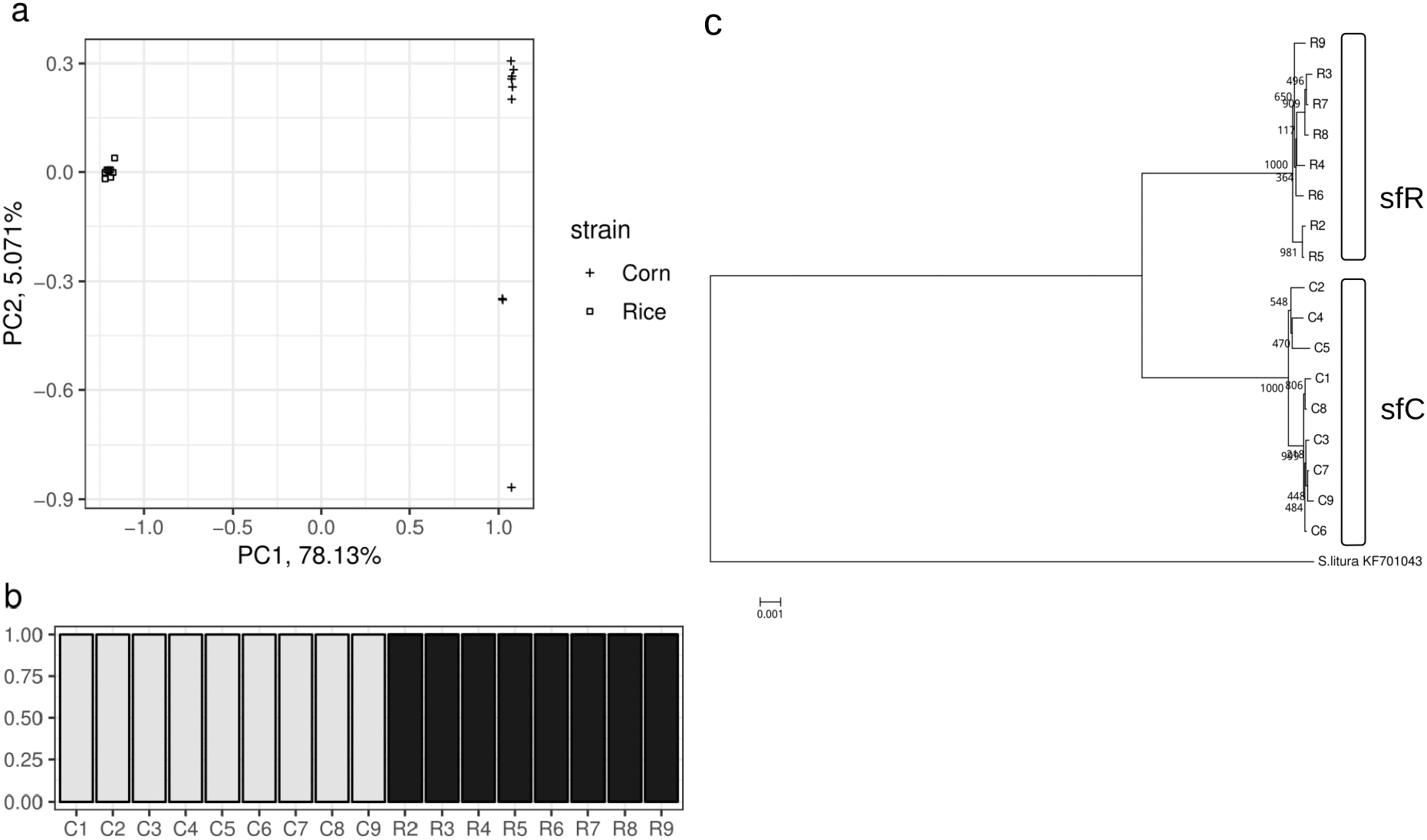
Mitochondrial genetic relationship between corn and rice strains. a) The result from principal component analysis. The red and blue circles represent individuals from sfC ad sfR, respectively. b) Ancestry coefficient results at K = 2. c) Phylogenetic tree reconstructed using minimum evolution approach.

A molecular clock study shows that the mitochondrial genomes diverged between sfC and sfR two million years ago(Kergoat et al. 2012), which corresponds to 2 × 10^7^ generations according to the observation from our insectarium (10 generations per year). Assuming that the *Ne* is 4 × 10^6^ for both strains, the number of generations during this mitochondrial divergence time is five times of *Ne*. We performed a simple forward simulation(Haller and Messer 2017) with a wide range of migration rate to test this divergence time can explain the level of observed genetic differentiation (Fst = 0.017). No simulation generates Fst equal or lower than 0.017 (Supplementary Figure 8), supporting that mitochondrial genomes diverged more anciently than nuclear genomes.

We investigated the role of the rest of outliers, denoted by genomic islands of differentiation in this paper. Genomic islands of differentiation have much lower π than the genomic average in both strains (Supplementary Figure 9), and sfC has a lower π than sfR (p = 0.0007; Wilcoxon rank sum test). This result suggests that the genomic islands of differentiation were targeted by linked selection, as a form of selective sweeps(Smith and Haigh 1974) or background selection(Charlesworth 2012), with a greater extent in sfC. dXY calculated from genomic islands of differentiation is on average lower than the genomic average (Supplementary Figure 10), suggesting that these sequences were targeted by linked selection after the split between sfC and sfR. PCA from genomic islands of divergence and genomic islands of differentiation shows that these two types of genomic islands have a clear grouping according to strains (Figure 5), which was observed from mitochondrial genomes (Figure 4a) but not from nuclear genomes (Figure 2a). Interestingly, the sequences of genomic islands of divergence have comparable genetic variability between sfC and sfR, whereas sfC has a lower genetic variability in the sequence of genomic islands of differentiation than sfR. From these results, we concluded that the sfC diverged from sfR by linked selection.

**Figure 5.**
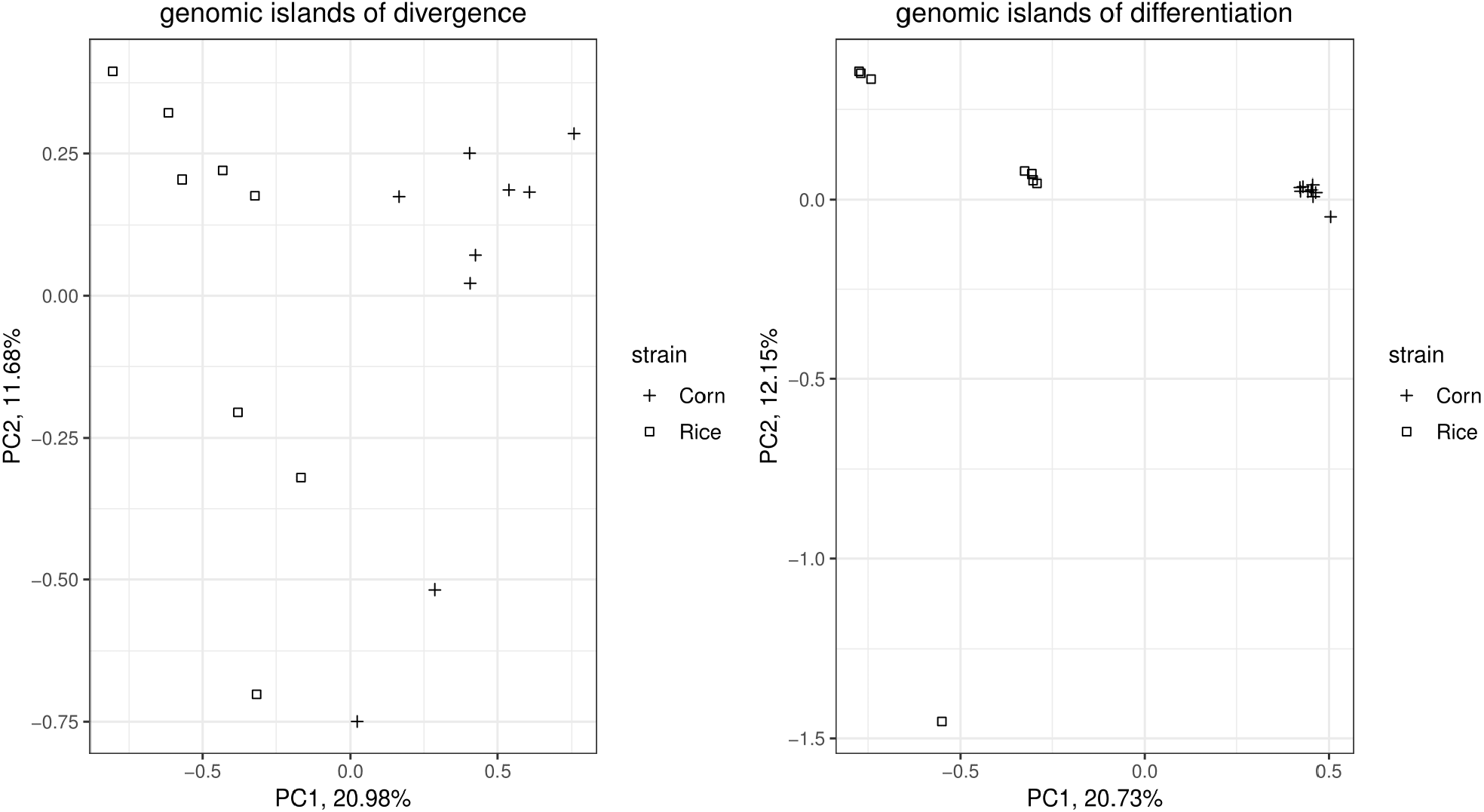
Genetic relationship among individuals in outliers of genetic differentiation. The result of principal component analysis from genomic islands of divergence (left), which have higher level of both relative level of genetic differentiation (hapflk score) and absolute level of genetic divergence (dXY), and genomic islands of differentiation (left), which have higher level of genetic differentiation (hapflk score) only.

We investigated the role of physical linkage by performing PCA with varying distances to the nearest genomic island of differentiation. When the distance is less than 1kb, genetic variations of sfC individuals are included within the range of genetic variation of sfR individuals (PC1 of the leftmost panel at Figure 6), while divergence of sfC from sfR is also supported (PC2 of the leftmost panel at Figure 6). If the distance is higher than 1kb, the divergence of sfC from sfR is not observed (Figure 6), suggesting that the effect of physical linkage to genomic islands of differentiation disappears rapidly as the distance increases. The short linkage disequilibrium in a species with large *Ne* is expected from a theoretical analysis(Feder and Nosil 2010) and reported from empirical cases(Sved et al. 2013; Song et al. 2015). These results show that physical linkages among targets of linked selection are not the primary cause of genomic differentiation.

**Figure 6.**
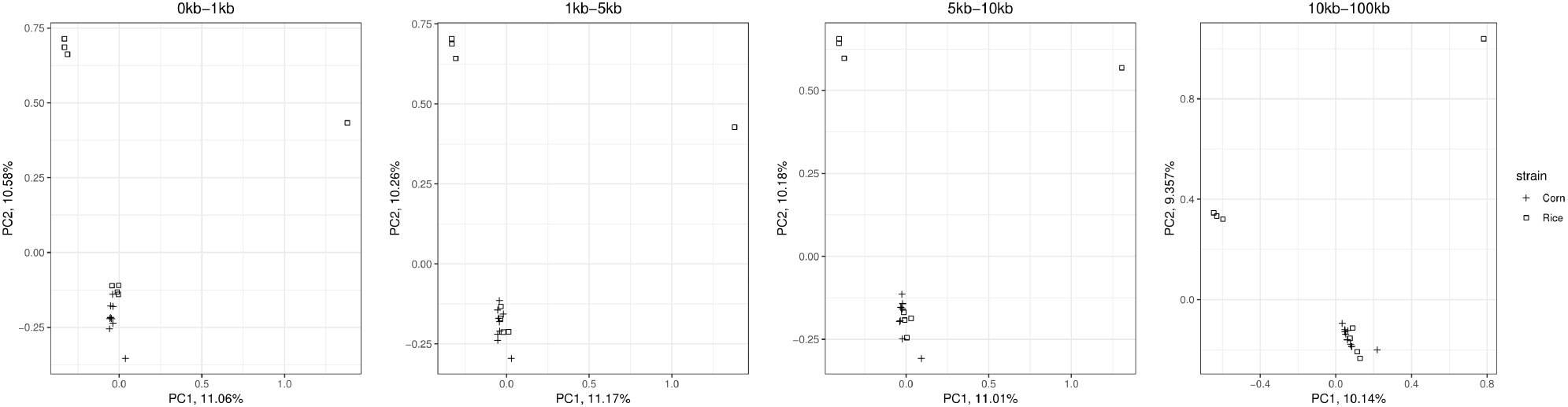
The effect of physical linkage to the genomic islands of genetic differentiation. The result of principal component analysis at varying distances from the nearest the genomic islands of genetic differentiation. The result is based on the mappings against refC. See Supplementary Figure 20 for the result based on the mapping against refR.

Then, we tested a possibility of genome hitchhiking(Feder and Nosil 2010; Feder, Gejji, et al. 2012), a hypothesis stating that genomic differentiation is caused by a genome-wide reduction in migration rate due to many loci under selection. If the strength of selection determines the level of genetic differentiation, a positive correlation between Fst and the strength of selection is expected. Alternatively, if a genomic reduction in migration rates dominates the effect of selection, this correlation is not expected. We assume that the exon density is a proxy for the strength of selection. Exon densities calculated in 100kb window are negatively correlated with π (Spearman’s ρ = −0.211, p < 2.2 × 10^−16^) (Figure 7), showing that the local genetic diversity pattern is affected by selection. Fst, however, is not significantly correlated with exon density (ρ = 0.021, p = 0.2032) (Figure 7). This result supports the hypothesis that a genomic reduction in migration rate dominates the variation of genetic differentiation due to selection.

**Figure 7.**
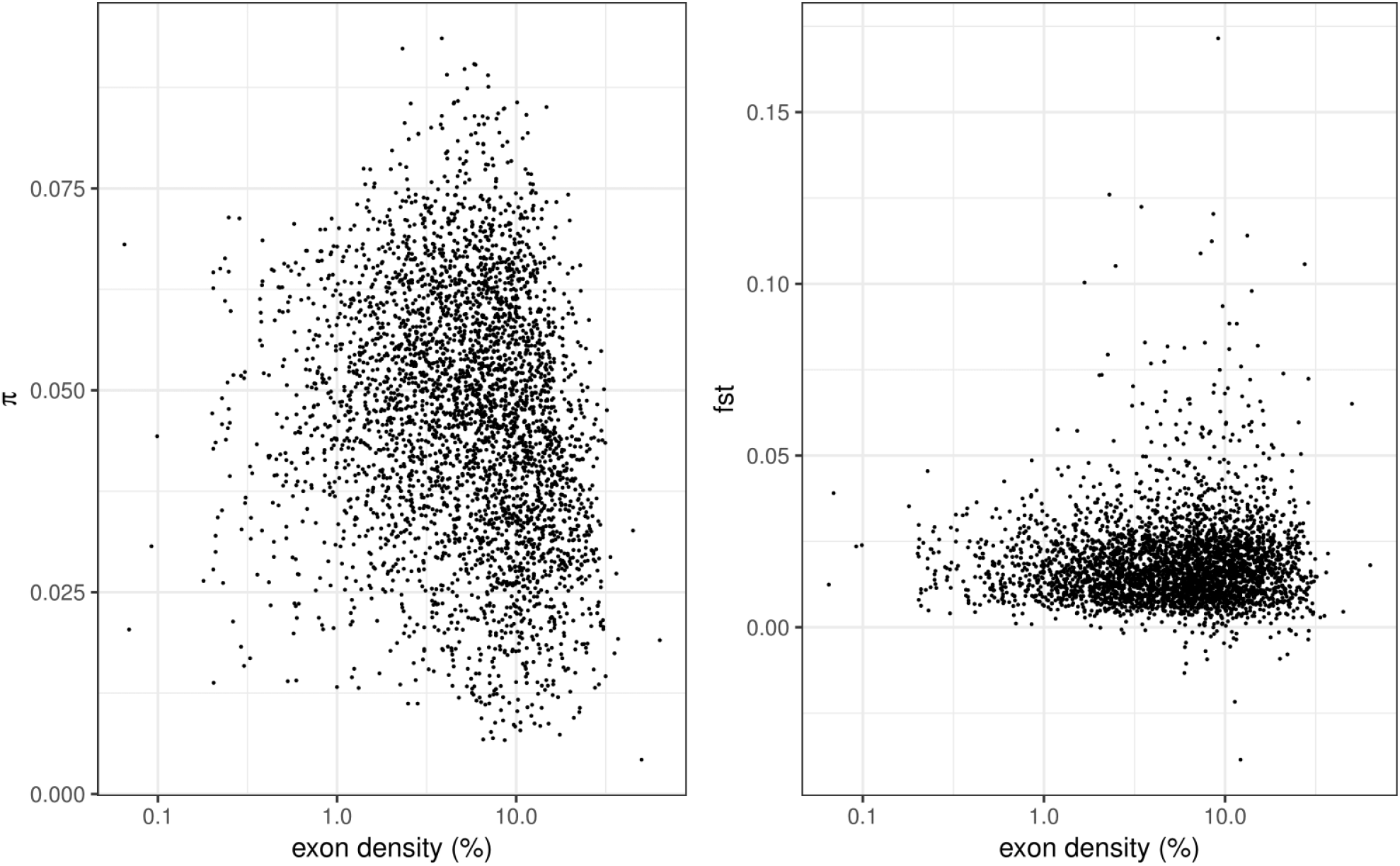
The effect of selection on local variation of diversity and differentiation. Plots showing the correlation of exon density with π (left) and Fst (right) calculated from 100kb windows, based on the mapping against refC. See Supplementary Figure 21 for the result based on the mapping against refR.

In principle, both selective sweeps and background selection may target these genomic islands of differentiation as linked selection. Background selection may cause genetic differentiation between populations only if these two populations are *a priori* differentiated by a geographical separation or a tight physical linkage to a target of selective sweeps. As sfC and sfR are sympatrically observed and the physical linkage among genomic islands of differentiation is not supported, as shown above, we assume that selective sweeps are mainly responsible for the genomic islands of differentiation and traits under adaptive evolution were inferred from the function of genes within genomic islands of differentiation. These islands contain 275 and 295 protein-coding genes from refC and refR, respectively (the full list can be found from Supplementary Table 4-5). These protein-coding sequences include a wide range of genes important for the interaction with host-plants, such as P450, chemosensory genes, esterase, immunity gene, and oxidative stress genes(Gouin et al. 2017) (Supplementary Table 3), suggesting that ecological divergent selection is important for genomic differentiation. Interestingly, cyc gene, which plays a key role in circadian clock(Rutila et al. 1998), is also included in the list of the potentially adaptively evolved genes. Thus, divergent selection on cyc might be responsible for pre-mating reproductive isolation due to allochronic mating time(Schöfl et al. 2009; Hänniger et al. 2017).

A QTL study shows that genetic variations in vrille gene can explain differentiated allochronic mating behavior in *S. frugiperda*(Hänniger et al. 2017). This gene is not found in the outliers. Fst calculated from a 10kb window containing this gene is 0.017 and 0.016 for refC and refR, respectively, which is similar to genomic average (0.017). Thus, it appears that this gene does not have a direct contribution to genomic differentiation.

## DISCUSSION

In this study, we showed that genetic differentiation between strains in *S. frugiperda* is initiated by the divergence of genes associated with female fecundity from the gene list in the genomic islands of divergence (Figure 8 to see a possible evolutionary scenario of genetic differentiation between sfC and sfR). Afterward, divergent selection targeting many loci appears to reduce the genome-wide migration between strains, which have low but significant genome-wide differentiation, in line with the genome hitchhiking model(Feder and Nosil 2010; Feder, Gejji, et al. 2012). The physical linkage among targets of linked selection appears to be unimportant for genomic differentiation in *S. frugiperda*. We observed that genomic islands of differentiation contain genes associated with interaction with host-plants. Thus, the adaptive evolution of this ecological trait appears to promote genomic differentiation between strains. A circadian gene (cyc) is also found from a genomic island of differentiation, and it is unclear whether this gene is associated with the assortative mating due to allochronic mating patterns in *S. frugiperda*. If genetic differentiation of this gene causes assortative mating, both divergent selection and assortative mating generate genomic differentiation by a genomic reduction in migration rate between strains, since assortative mating generates the same footprints on DNA sequences as divergent selection. In short, the genetic differentiation was initiated by disruptive selection on traits associated with female fecundity in *S. frugiperda*, and divergent selection targeting on many loci enables the transition from genetic to genomic differentiation without the involvement of physical linkages among targets or chromosomal rearrangements.

**Figure 8.**
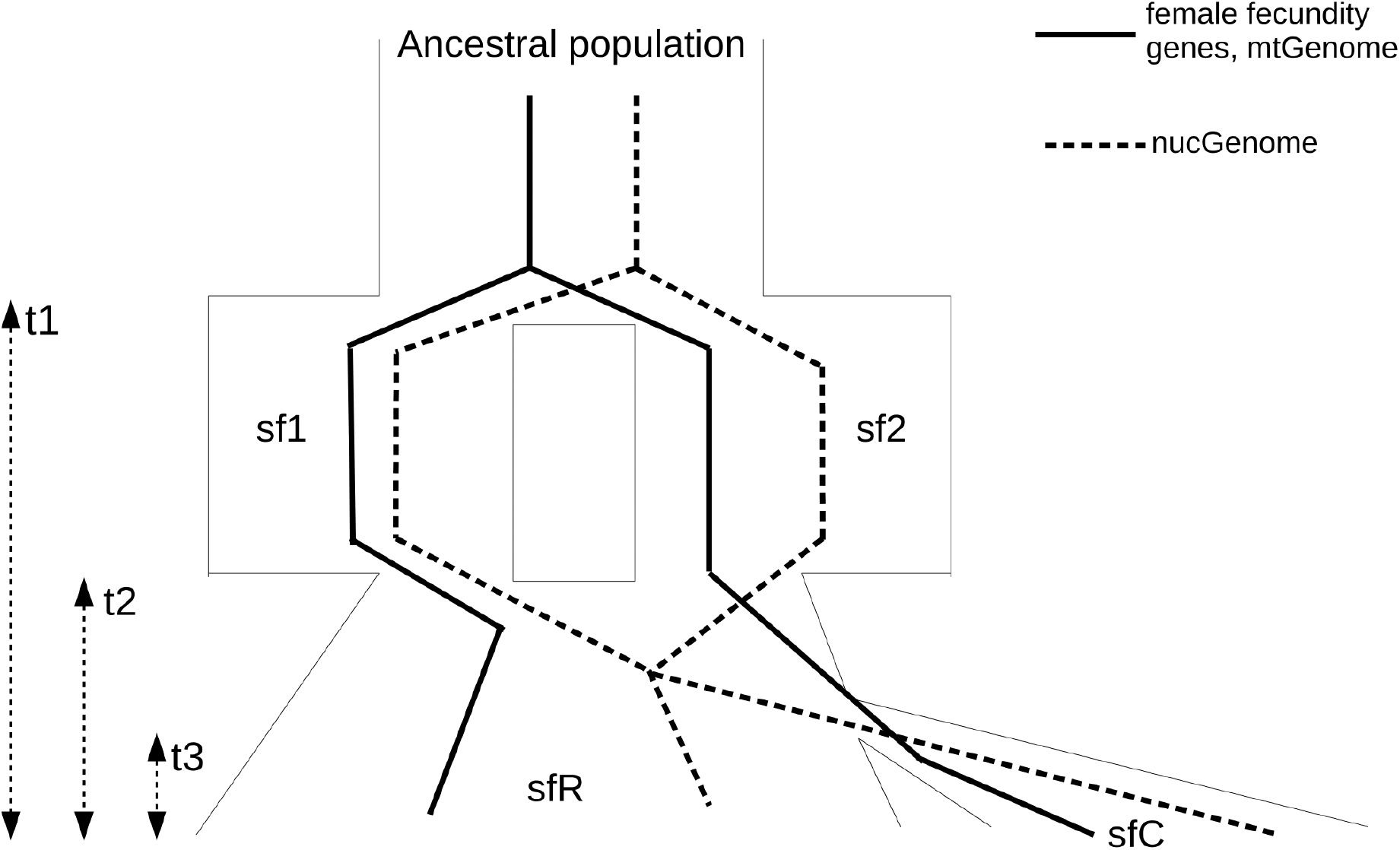
A possible evolutionary scenario of genetic differentiation. The average genealogy of mitochondrial genomes, female fecundity genes (solid lines), and nuclear genomes (dashed lines) are depicted. In this scenario, an ancestral population was split into two populations, sf1 and sf2, at t1. At t2, two populations were merged by hybridization, and extant sfR was generated. However, local gene flow between sf1 and sf2 was inhibited at female fecundity genes because hybrids of these genes had a reduction in fitness. Thus, the genealogy of the female fecundity genes remained separated, and sequences were kept diverging. The genealogy of mitochondrial genomes is the same with the female fecundity genes because of selection on females and maternal inheritance. After t3, divergent selection targeting many genes caused a genetic differentiation according to the sequences of mitochondrial genomes and female fecundity genes by reducing genomic migration rate, and extant sfC was generated.

The heterozygosity of these strains is unprecedented high, as the calculated π is 0.043-0.044. In two other Noctuid pests, *S. litura* and *Helicoverpa armigera*, π calculated from multiple populations across their distribution area ranges from 0.0019 to 0.016(Cheng et al. 2017), and from 0.008 to 0.01(Anderson et al. 2018), respectively. *Heliconius melpomene*, a butterfly species, has π between 0.021 and 0.029(Martin et al. 2016). To explain the extremely high level of heterozygosity in *S.frigiperda*, we first checked the possibility that a considerable proportion of identified variants is false positives. We performed additional filterings, on the top of applied ones, by including additional 12 criteria. These additional filterings discarded only 34 out of 48,981,416 and 17 out of 49,832,320 variants from the mapping against refC and refR, respectively. Thus, we exclude the possibility that false positives caused the high level of heterozygosity. We inferred past demographic history using pairwise sequentially Markovian coalescent(Li and Durbin 2011) based on assumptions that generation time is the same with lab strains at our insectarium (10 generation/yr) and mutation rate is the same with *H.melpomene* (2.9 × 10^−9^/site/generation)(Keightley et al. 2015). Extremely rapid population expansions were inferred from both two strains (*Ne* was increased from 9.6 × 10^5^ to 1.2 × 10^7^) between 10 mya and 100 mya (Supplementary Figure 11). A possible explanation of this rapid expansion is the merge of genetically diverged ancestral populations by hybridization. In this scenario (Figure 8), two populations were separated by geographical barriers and genetically differentiated. At some moment, the geographical barriers were removed, and these populations started to be merged by hybridization. As the merged population maintains a large proportion of variants, this population has a high level of heterozygosity. This population is extant sfR. Afterward, a group of sfR started to diverge by ecological divergent selection and assortative mating, and this group became the extant sfC. This process of genomic differentiation is similar to the description of a speciation process in cichlid (Meier et al. 2018), but we proposed that this process may occur even among populations in single species. This explanation does not exclude the possibility of direct selection on mitochondrial genes(Orsucci et al. 2018).

The pattern of genomic differentiation can be different among different geographical populations. For example, pairs of different geographical populations may have different levels of genomic differentiation (Fst). The genomic islands of differentiation can also be different if a proportion of divergent selection is specific to a single geographical population. Therefore, it is worthwhile to test if the same loci are identified as genomic islands of divergence across diverse geographic populations. If levels of genomic differentiation vary among different geographical populations in *S. frugiperda*, it might be possible to find a pair of strains that enter to a phase of genome-wide congealing. Attempts to find the process towards complete genomic differentiation often termed ‘speciation continuum’ are typically based on closely related multiple species(Martin et al. 2013; Riesch et al. 2017). However, different species may have experienced very different evolutionary histories. Thus, studying a single species with varying levels of genetic differentiation might shed light on the exact process of genomic differentiation.

Several genetic markers have been proposed to identify strains, including mitochondrial CO1(Pashley 1989), sex chromosome FR elements (Lu et al. 1994), and Z-linked TPI(Nagoshi 2010). We found that FR elements are a reliable marker to identify strains (Supplementary Figure 12). TPI is included in the gene list within the genomic island of differentiation, and dXY from TPI (0.0345) is slightly lower than genomic average (mean is 0.0384 with 0.0383-0.0386 of 95% confidence interval). Thus, the genetic differentiation of TPI appears to occur after the initiation of genetic differentiation between sfC and sfR. The concordance of identified strains between mitochondrial CO1 and TPI can be as low as 74% (Table 5 at (Nagoshi 2010)), and this imperfect concordance might be due to the different divergence time. Thus, we propose to use mitochondrial markers to identify strains for unambiguous strain identification.

The process of speciation proposed in this study can be further tested based on insect rearing or lab experiments (such as CRISPR/CAS9). For example, we proposed in this study that female fecundity could be a key trait that initiated genetic differentiation between strains because genes associated with this trait appears to have a resistance against gene flow. The reason for this resistance can be a reduction in hybrid fitness, and we can test this possibility by insect-rearing. We also raise a possibility in this paper that cyc gene might be associated with allochronic mating behavior, and we can test this possibility using CRISPR/CAS9 experiment as well. These future studies will shed light on the relationship between genotypes and phenotypes that plays critical roles in the process of speciation.

## MATERIALS AND METHODS

We extracted high molecular weight DNA using MagAttract© HMW kit (Qiagen) from one pupa of sfC and two pupae of sfR with a modification of the original protocol to increase the yield. The quality of extraction was assessed by checking DNA length (> 50kb) on 0.7% agarose gel electrophoresis, as well as pulsed-field electrophoresis using the Rotaphor (Biometra) and gel containing 0.75% agarose in 1X Loening buffer, run for 21 hours at 10°C with an angle range from 120 to 110° and a voltage range from 130 to 90V. DNA concentration was estimated by fluorimetry using the QuantiFluor Kit (Promega), 9.6 μg and 8.7 μg of DNA from sfC and sfR, respectively, which was used to prepare libraries for sequencing. Single-Molecule-Real-Time sequencing was performed using a PacBio RSII (Pacific Biosciences) with 12 SMRT cells per strain (P6-C4 chemistry) at the genomic platform Get-PlaGe, Toulouse, France (https://get.genotoul.fr/). The total throughput is 11,017,798,575bp in 1,513,346 reads and 13,259,782,164bp in 1,692,240 reads for sfC and sfR, respectively. The average read lengths are 7,280bp and 7,836bp for sfC and sfR, respectively.

We generated assemblies from Illumina paired-end sequences(Gouin et al. 2017) (166X and 308 X coverage for sfC and sfR, respectively) using platanus(Kajitani et al. 2014). Then, errors in PacBio were corrected using Ectools(Gurtowski 2017), and uncorrected reads were discarded. The remaining reads are 8,918,141,742bp and 11,005,855,683bp for sfC and sfR, respectively. The error-corrected reads were used to assemble genome sequences using SMARTdenovo(Ruan 2017). The paired-end Illumina reads were mapped against the genome assemblies using bowtie2(Langmead and Salzberg 2012: 2), and corresponding bam files were generated. We improved the genome assemblies with these bam files using pilon(Walker et al. 2014). For the genome assemblies of sfC, both Illumina paired-end and mate-pair reads were mapped the genome assemblies using bwa(Li and Durbin 2010), and scaffolding was performed using BESST(Sahlin et al. 2016). Since only paired-end libraries were generated from sfR in our previous study(Gouin et al. 2017), we used only paired-end sequences to perform scaffolding for sfR. The gaps were filled using PB-Jelly(Rizk et al. 2014). The correctness of assemblies was assessed using insect BUSCO (insecta_odb9)(Simão et al. 2015). Then, protein-coding genes were annotated from the genome sequences using MAKER(Cantarel et al. 2008). First, repetitive elements were masked using RepeatMasker(RepeatMasker). Second, *ab initio* gene prediction was performed with protein-coding sequences from two strains in *S. frugiperda*(Gouin et al. 2017) and *Helicoverpa armigera* (Harm_1.0, NCBI ID: GCF_002156995), as well as insect protein sequences from *Drosophila melanogaster* (BDGP6) and three Lepidoptera species, *Bombyx mori* (ASM15162v1), *Melitaea cinxia* (MelCinx1.0), and *Danaus plexippus* (Dpv3) in ensemble metazoa. For transcriptome sequences, we used reference transcriptome for sfC(Legeai et al. 2014) and locally assembled transcriptome from RNA-Seq data from 11 samples using Trinity(Grabherr et al. 2011) for sfR. Third, two gene predictors, SNAP(Korf 2004) and Augustus(Stanke et al. 2006), were trained to improve gene annotations. Multiple trainings of the gene predictors do not decrease Annotation Edit Distance Score. Thus, we used the gene annotation with only one training. Fourth, we discarded all gene prediction if eAED score is greater than 0.5.

Paired-end Illumina resequencing data from nine individuals from each of corn and rice strains in *S. frugiperda* is used to identify variants. Low-quality nucleotides (Phred score < 20) and adapter sequences in the reads were removed using AdapterRemoval(Schubert et al. 2016). Then, reads were mapped against reference genomes using bowtie2, with very exhaustive local search parameters (-D 25 -R 5 -N 0 -L 20 -i S,1,0.50), which is more exhaustive search than the –very-sensitive parameter preset. Potential PCR or optical duplicates were removed using Picard tool(Picard 2018). Variants were called using samtools mpileup(Li et al. 2009) only from the mappings with Phred mapping score higher than 30. Then, we discarded all called positions unless a genotype is determined from all individuals and variant calling score is higher than 40. We also discarded variants if the read depth is higher than 3,200 or lower than 20.

We used vcftools to calculate population genetics statistics, such as π and Fst(Danecek et al. 2011). Watterson’s θ and d_XY_ were calculated using house-perl scripts. To estimate the genetic relationship among individuals, we first converted VCF files to plink format using vcftools, then PCA was performed using flashpca(Abraham et al. 2017). For ancestry coefficient analysis, we used sNMF(Frichot et al. 2014) with K values ranging from 2 to 10, and we chose the K value that generated the lowest cross entropy.

Phylogenetic tree of the nuclear genome was generated using AAF(Fan et al. 2015). As an outgroup, we used simulated fastq files from the reference genomes of *S. litura*(Cheng et al. 2017) using genReads(Stephens et al. 2016) with an error rate equal to 0.02. Reads were mapped against the mitochondrial genome (KM362176) using bowtie2(Langmead and Salzberg 2012: 2) to generate the mitochondrial phylogenetic tree, and variants were called using samtools mpileup(Li et al. 2009). From the mitochondrial VCF file, a multiple sequence alignment was generated using house-perl script. Then, the whole mitochondrial genome from *S. litura* (KF701043) was added to this multiple sequence alignment, and a new alignment was generated using prank(Löytynoja 2014). The phylogenetic tree was reconstructed from this new alignment using FastME(Lefort et al. 2015) with 1,000 bootstrapping.

The outliers of genetic differentiation were identified from hapFLK scores calculated from hapflk software(Fariello et al. 2013). As the computation was not feasible with the whole genome sequences, we randomly divided sequences in the genome assemblies into eight groups. Fst distributions from these eight groups were highly similar between each other (Supplementary Figure 13). P-values showing the statistical significance of genetic differentiation were calculated from each position using scaling_chi2_hapflk.py in the same software package.

The reference genome and gene annotation are available from BioInformatics Platform for Agroecosystem Arthropods together with the genome browser (https://bipaa.genouest.org/is/). This data can be found at European Nucleotide Archive (https://www.ebi.ac.uk/ena) as well (project id: PRJEB29161). Resequencing data is available from NCBI Sequence Read Archive, and corresponding project ID is PRJNA494340.

## ACKNOWLEDGMENTS AND FUNDING INFORMATION

This work was supported by a grant from the department of Santé des Plantes et Environnement at Institut national de la recherche agronomique for K.N. (adaptivesv), by a grant from the French National Research Agency for E.A. (ANR-12-BSV7-0004-01), and by a grant from Institut Universitaire de France for N.N.

## AUTHOR CONTRIBUTIONS

K.N. and N.N. designed the study; K.N performed the genome assembling and the analyses; S.N. and E.A. performed SMRT sequencing; S.R. and A.B. performed gene annotation; K.N. wrote the manuscript.

